# Single-Cell Reprogramming of Mouse Embryo Development Through a Critical Transition State

**DOI:** 10.1101/140913

**Authors:** Masa Tsuchiya, Alessandro Giuliani, Kenichi Yoshikawa

## Abstract

Our work dealing with the temporal development of the genome-expression profile in single-cell mouse early embryo indicated that reprogramming occurs via a critical transition state, where the critical-regulation pattern of the zygote state disappears. In this report, we unveil the detailed mechanism of how the dynamic interaction of thermodynamic states (critical states) enables the genome system to pass through the critical transition state to achieve genome reprogramming.

Self-organized criticality (SOC) control of overall expression provides a snapshot of self-organization and explains the coexistence of critical states at a certain experimental time point. The time-development of self-organization is dynamically modulated by exchanges in expression flux between critical states through the cell nucleus milieu, where sequential global perturbations involving activation-inhibition of multiple critical states occur from the early state to the late 2-cell state. Two cyclic fluxes act as feedback flow and generate critical-state coherent oscillatory dynamics. Dynamic perturbation of these cyclic flows due to vivid activation of the ensemble of low-variance expression (sub-critical state) genes allows the genome system to overcome a transition state during reprogramming.

Our findings imply that a universal mechanism of long-term global RNA oscillation underlies autonomous SOC control, and the critical gene ensemble at a critical point (CP) drives genome reprogramming. Unveiling the corresponding molecular players will be essential to understand single-cell reprogramming.

## Introduction

In mammalian embryo development, a large number of molecular-level epigenetic studies [1-3] revealed the occurrence of stunning global epigenetic modifications on chromatins (DNA + histones) associated with reprogramming processes in mammalian embryo development. However, the genome-wide principle that drives such extremely complex epigenetic modifications is still unknown.

In our previous studies, based upon transcriptome experimental data for seven distinct cell fates [4], we recognized that a self-organized critical transition (SOC) in whole-genome expression plays an essential role in the change of the genome expression state at both the population and single-cell levels (see **Methods,** for more details [4-6]).

Essential points of SOC control of overall expression can be summarized as: i) SOC control of overall expression represents self-organization of the coexisting critical states (distinct response expression domains) through a critical transition. Temporal variance expression (normalized root mean square root fluctuation: *nrmsf*; see **Methods**) acts as an order parameter in self-organization. Distinct critical states can be observed in a transitional behavior of ensemble expression profile (e.g., bimodality coefficient) or bifurcation of ensemble expression state (e.g., probability density profile) according to *nrmsf*. Coherent behaviors emerge from stochastic expression within critical states (coherent-stochastic behaviors; see non-equilibrium statistical mechanism underpinning the spontaneous emergence of order out of disorder [7]) as the collective behaviors of groups with more than around 50 genes (mean-field approach) [6,8,9].

ii) Self-organization based on SOC occurs through distinguished critical behaviors: *sandpile-type critical behavior (criticality)* and *scaling-divergent behavior.* In sandpile-type criticality, the divergence of two different regulatory behaviors occurs at a critical point (summit of sandpile) neatly separating up- and down-regulation. Sandpile-type criticality emerges spontaneously by grouping of different gene expression according to their fold change. Reprogramming or cell-fate change is observed as erasure of criticality of an initial-state (such as an early embryo state). Scaling-divergent behavior emerge on averaging behavior of gene groups expression according to *nrmsf*, producing both linear (scaling) and divergent domains in a log-log plot. This divergent behavior anticipates a transition [10]. Since *nrmsf* is an order parameter, the above separation of scaling domains (**Figure 1**) shows how self-organization occurs in overall expression and it is not confined to few ‘master genes’.

**Figure 1:**
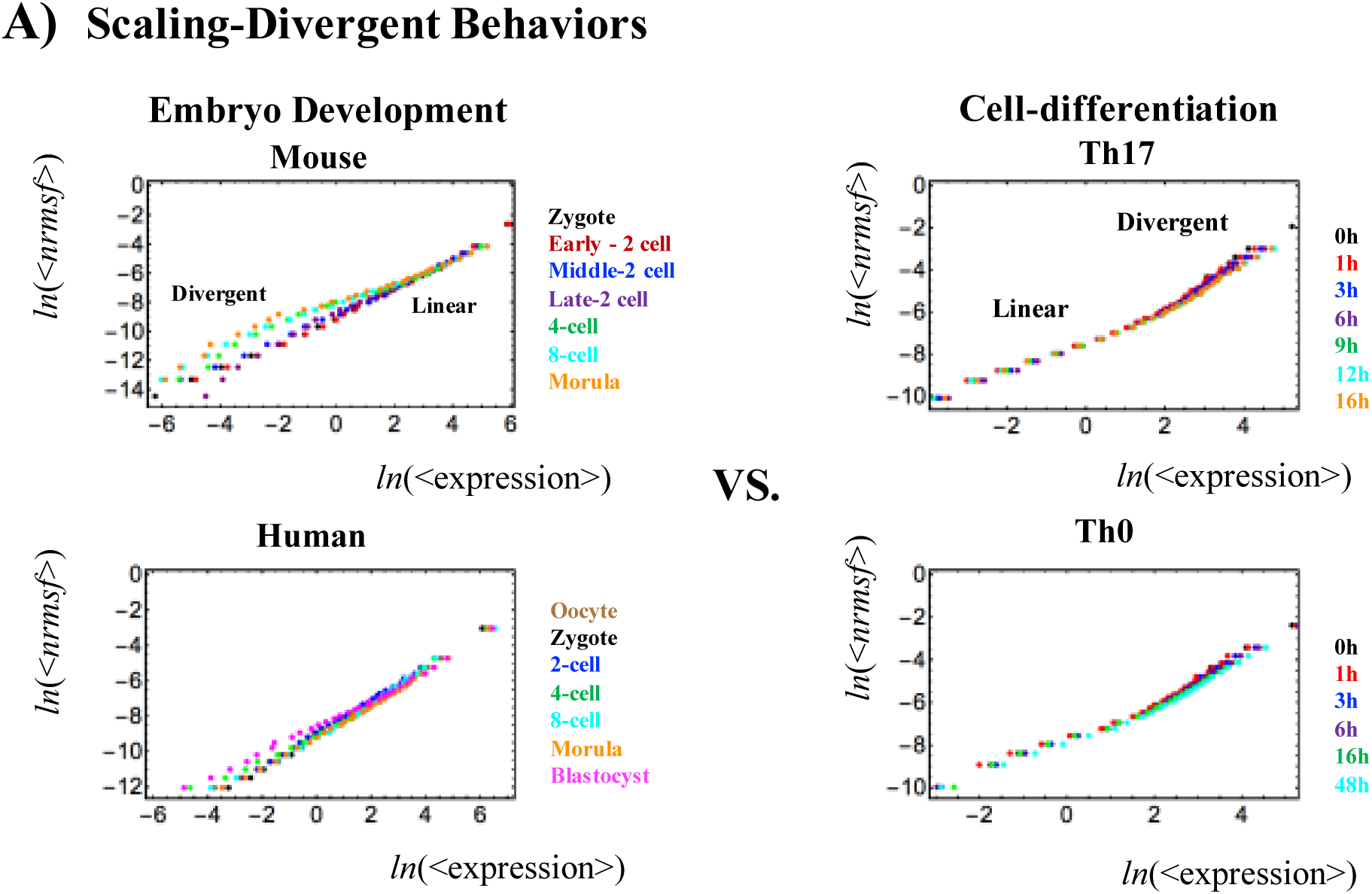
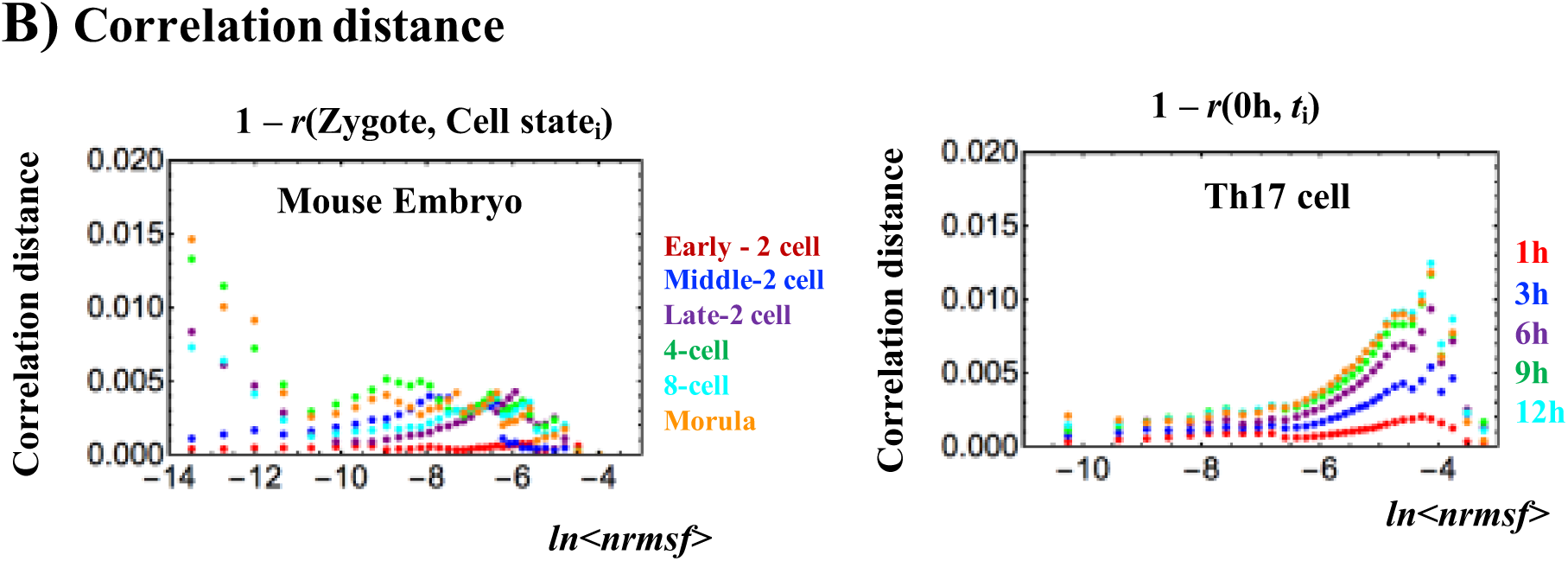
Opposite genome avalanche behaviors in early embryo development and immune cell differentiation: **A):** Genome avalanches, scaling-divergent behaviors in overall expression, as important features of the SOC control of overall expression are evident in the log-log plot of averaging behaviors: (left panel: mouse (first row) and human (second) embryo development; right panel; Th17 (first) and Th0 (second) terminal cell differentiation). Log-log plots represent the natural logarithm of the group average (< >) of expression (x-axis) and nrmsf (y-axis) (n = 685 (mouse), 666 (human), 525 (Th17) and 504 RNAs (Th0) for each dot), where overall expression is sorted and grouped (25 groups) according to the degree of nrmsf (normalized root mean square root fluctuation: see Methods). Two distinct biological processes (reprogramming in early embryo development versus immune cell differentiation) show opposite scaling-divergent behaviors. Scaling behavior occurs in the ensemble of high-variance RNA expression (high region of nrmsf: super-critical state; see **Figure 3A**) in early embryo development and divergent behavior in the ensemble of low-variance RNA expression (low region of nrmsf: sub-critical state), whereas the T cell terminal cell fate (single cell) has opposite behaviors. **B): Correlation distances**: It is expressed as (1 - r), where r is the Pearson correlation coefficient between the zygote and another development states gene expression profiles. This distance corresponds to the relative change in expression profile on the whole genome scale and it is computed separately for different critical states. The result supports the difference in scaling-divergent behaviors between embryo development and cell differentiation: in mouse embryo, correlation behaviors significantly change in low nrmsf, whereas in terminal Th17 cell, significant change occurs in high nrmsf.

SOC control of overall expression can be seen in an emergent level of collective behaviors. The presence of a ‘collective organization layer’ in gene expression, while a naturally encompassed by the existence of ‘tissue specific gene expression profiles’ across the whole genome, seems at odds with the major part of biological experimentation strictly focusing on gene-specific rules. The opposition emicroscopic rules versus ecollective behavior’, is a classical issue in science: it is worth stressing the presence of microscopic laws does not falsify the statistical approach (that in this condition could be considered as a surrogate of a more efundamental microscopic approach) while the statistical eemergent way can hold even in absence of microscopic deterministic rules [11]. In this last case, the efundamental level corresponds to the collective organization layer [11, 12]. As for our specific case, a gene-by-gene set of deterministic microscopic rules affecting thousands of gene products (some of which have very low concentration inside single cells and thus, undergoing high stochastic fluctuation) is expected to be both more unreliable and energy consuming than self-organized criticality that in turn is very clearly emerging as different scaling regimens along order parameter (**Figure 1)**

**Figure 2** reports the above sketched phenomena along the time arrow, clarifying the opposite behavior of early embryo development (going toward the erasure of the previous organization pattern relative to zygote) and terminal differentiation (going toward the establishment of a stable organization pattern).

**Figure 2:**
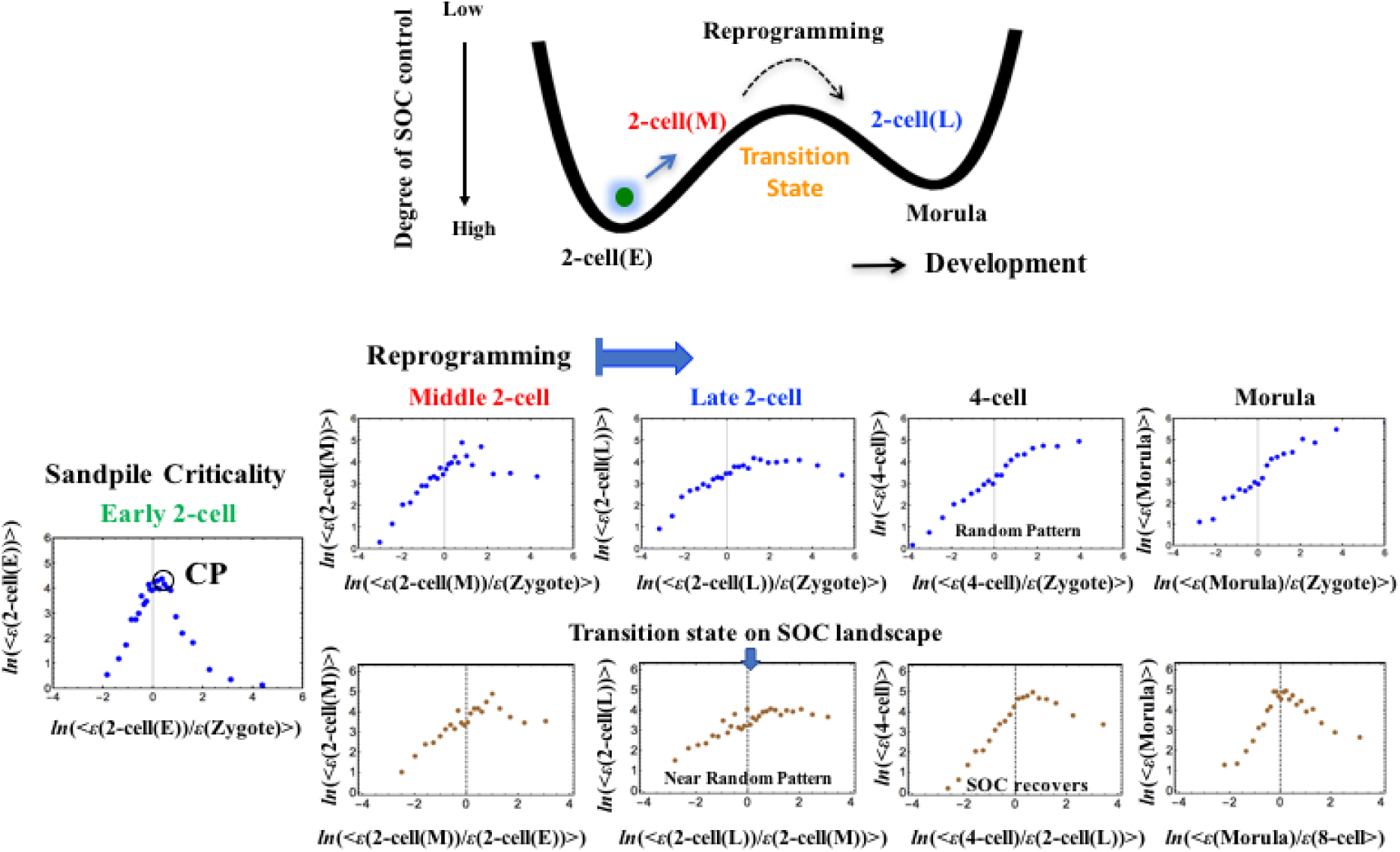
Timing of genome-state change on the SOC-control landscape through a transition state in a single cell: The timing of the genome-state change was revealed in the erasure of sandpile-type criticality of the zygote state. The erasure of zygote criticality (second row) occurs after the middle 2-cell state to reveal a stochastic expression pattern as a linear correlative behavior (refer to S2 Fig. in [4]). The transition of SOC control through non-SOC control (stochastic pattern) shows an SOC-control landscape: a valley (SOC control: zygote - early 2-cell states) - ridge (non SOC control: middle - late 2-cell states) - valley (SOC control: 8-cell - morula states); a high degree of SOC control for a well-developed shape of the sandpile-type criticality, an intermediate degree of SOC control for a weakened (broken) sandpile, and a low degree for non-SOC control. At the middle - later 2-cell states (third row), sandpile-type criticality disappears and recovers thereafter; this indicates the existence of a transition state in the SOC control landscape (see the detailed mechanism in **Figure 7**). Sandpile-type critical behaviors, which exhibit diverging up- and down-regulation at a critical point (CP), emerge in overall expression sorted and grouped according to the fold-change in expression (see stochastic resonance in [6]). Each group (net 25 groups) contains n = 685 RNAs. The x- and y-axes show the natural logarithm of the fold-change in each group average of RNA expression between the *s_i_* and *s_j_* states (*i*^th^ and *j*^th^ cell states; **Methods**) represented by ln(<*ε*(*s_i_*)/*ε*(*s_j_*)>) and of RNA expression (ln(<*ε*(*s_i_*)>)), respectively.

Recently, we demonstrated that essentially the same critical-state dynamics that we observed for cell differentiation processes [4,6] are also present in overall RNA expression in single-cell mouse embryo development, which is particularly relevant to give further proof of SOC control as a universal characteristic [13]. Overall RNA expression and its dynamics exhibit typical features (genome avalanche and sandpile type criticality) of self-organized criticality (SOC) control in mouse embryo development; **Figure 1** shows a genome avalanche, i.e., scaling-divergent behavior in a log-log-scale plot between expression and the temporal variance of expression (*nrmsf).* **Figure 2** shows that sandpile-type criticality (critical behavior) of the zygote state survives at the early 2-cell state and disappears after the middle 2-cell state to reach a stochastic pattern in the 4-cell state (linear pattern revealed in randomly shuffled overall expression: S2 Fig. in [4]).

Importantly, genome avalanches (**Figure 1**) reveal that the regions of scaling and divergent regions in terms of *nrmsf* are opposite between embryo development and cell differentiation at a single-cell level. Since *nrmsf* is an order parameter [4-6], this reveals the opposite order of self-organization; note that similar scaling-divergent order for cell differentiation is also observed at a cell-population level in cancer cells [4,6]. Furthermore, since chromosomes exhibit fractal organization [14,15], the power law behavior may reveal a quantitative relation between the aggregation state of chromatin through *nrmsf* and the average expression of an ensemble of genes, which is opposite in embryo development and cell differentiation, and the existence of coherent waves of condensation/de-condensation in chromatin in cell development, which is transmitted from low temporal expression region (low *nrmsf* region) in reprogramming of the embryo genome, to high temporal expression region (high *nrmsf*) in terminal somatic cell differentiation.

On the other hand, the erasure of sandpile-type criticality (**Figure 2**) indicates that i) the memory of early embryogenesis in the zygote is lost after the middle 2-cell state: a significant change in the genome-state, i.e., the occurrence of reprogramming, and ii) the SOC control landscape, showing valley (SOC control) - ridge (non SOC control) - valley (SOC control), exhibits a critical transition state at the middle - late 2-cell states through a stochastic pattern, in which SOC control (sandpile-type criticality) disappears (see more in [4] and [13]).

In this report, we investigate the self-organizing dynamics of whole RNA expression in mouse embryo development [16] to unveil how genome reprogramming occurs by passing through a critical transition state. This is elucidated by a dynamic expression flux analysis (quantitative evaluation of perturbation based on self-organization through SOC).

The results of the flux dynamics suggest that the critical gene ensemble of the critical point plays an essential role in reprogramming. Our results suggest that the SOC control mechanism of genome dynamics is rather universal among several distinct biological processes [4]. We think our findings may provide a universal classification scheme for phenomena with far-from-equilibrium phase transitions, which has been missing in past studies [17].

## Results

### I. Dynamic Interaction of Critical States

#### A) Sloppiness of Mouse RNA Expression Dynamics: Coherent-Stochastic Behaviors in Critical States

As has been demonstrated in cell differentiation [4-6], mouse embryo overall RNA expression is self-organized into distinct response domains (critical states). This can be seen as a transitional change in expression profile of groups of genes according to *nrmsf* (**Figure 3A**), which generates three different gene expression states: ‘super-critical’, ‘near critical’ and ‘sub-critical’ respectively.

**Figure 3:**
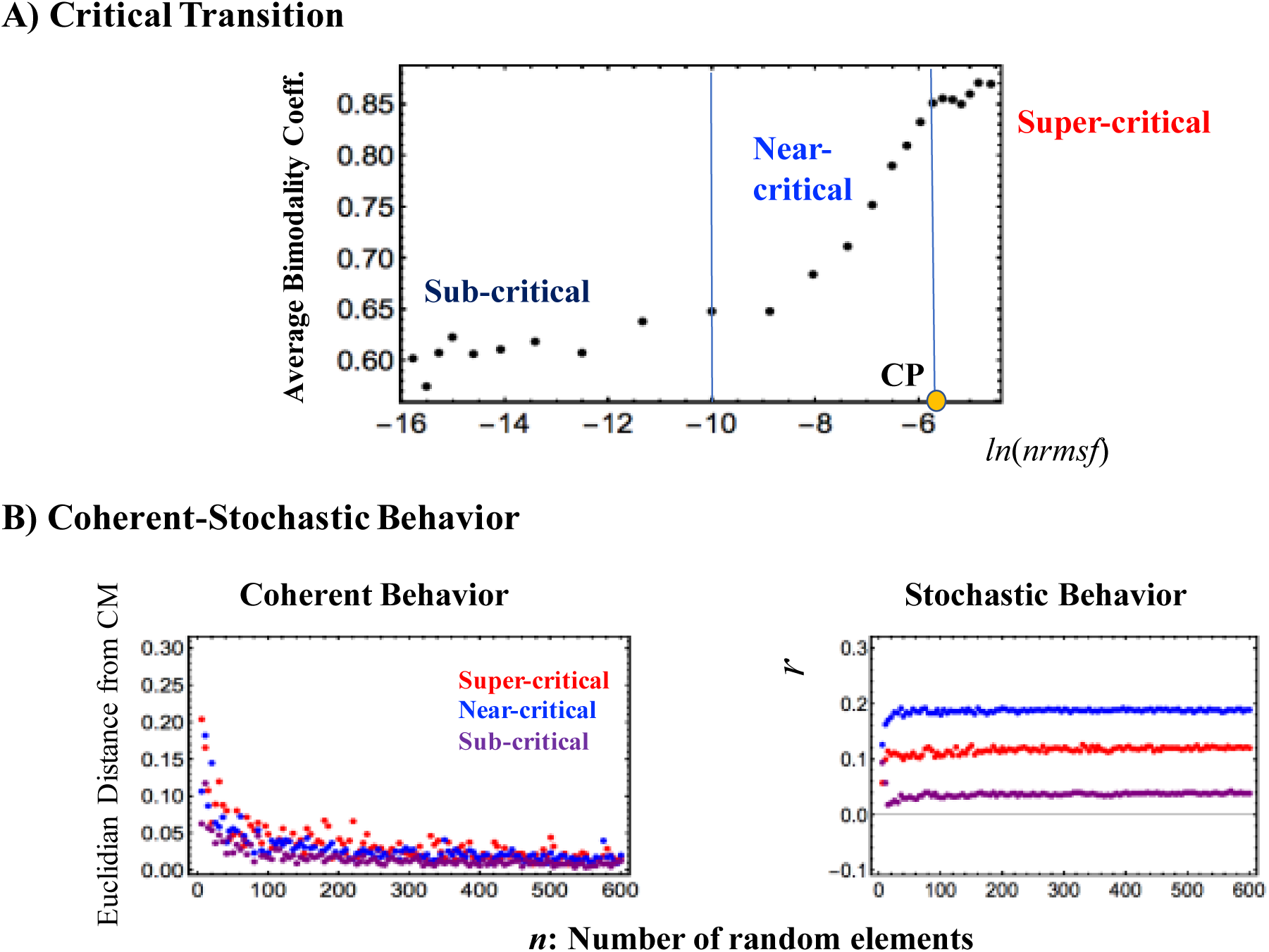
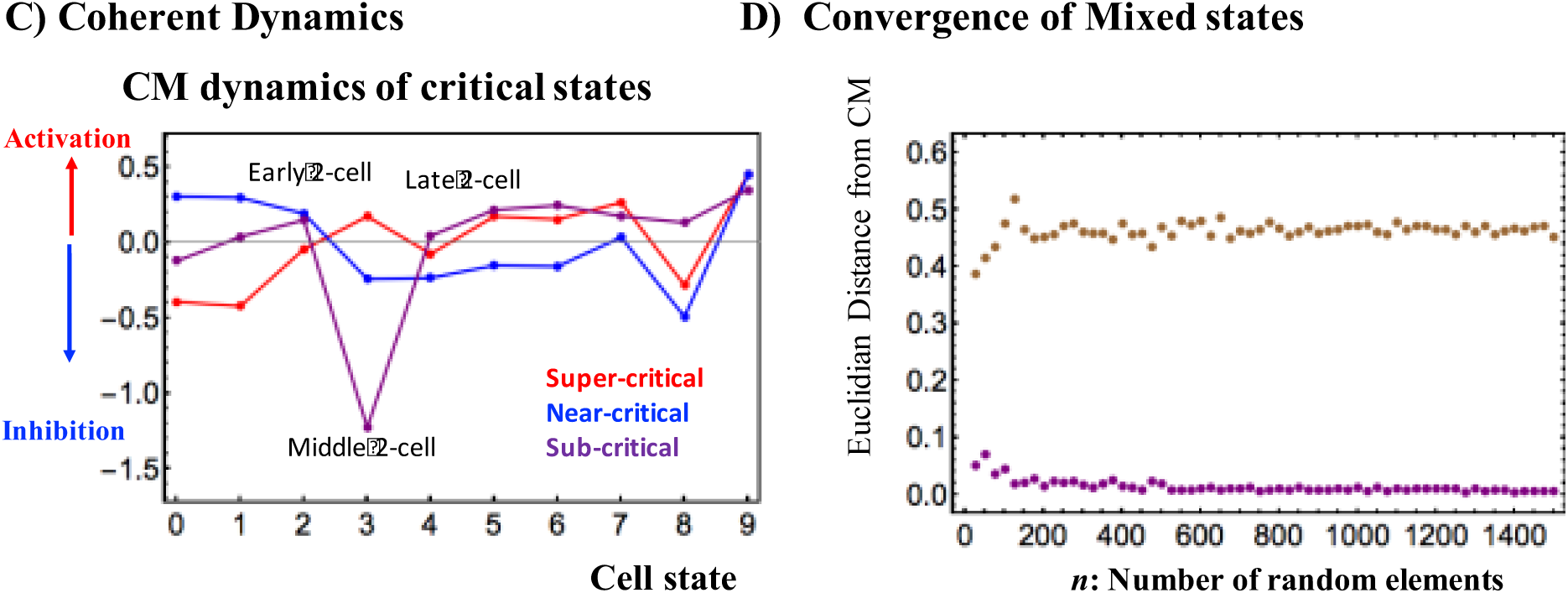
Critical states and their coherent stochastic behaviors: A) Estimation of the bimodality coefficients (averaged over all cell states) according to nrmsf. Overall RNA expression is sorted and grouped according to the degree of nrmsf. The nrmsf grouping is made at a given sequence of discrete values of nrmsf (*x_i_*: integer and integer + 0.5 from -15 to -4) with a fixed range (*x_i_* - 0.5 < *x_i_* < *x_i_* + 0.5), and the corresponding average of the Sarle’s bimodality coefficient (**Methods**) over the number of cell states is evaluated. The result exhibits a transitional behavior to distinguish (averaged) critical states with respect to the value of ln<*nrmsf*>: sub-critical state (N= 4913 RNAs)< -8.8; -8.8 <near-critical state (N = 9180)< -5.4; -5.4 <super-critical state (N= 3408), where < > indicates the ensemble average of the nrmsf grouping. A critical point (CP: summit of sandpile criticality) exists around the edge between the near- and super-critical states, where the CP exists around ln(<*nrmsf*>) ~ -5.5 (**Figure 2 and Figure 4**). B) Coherent stochastic behaviors are revealed: stochastic expression within a critical state (right panel) is confirmed by the low Pearson correlation of ensembles of RNA expression between samples randomly selected from a critical state (averaged over the number of cell states with 200 repeats). The degree of stochasticity further supports the existence of distinct critical states. In the left panel, the Euclidian distance (averaged over 200 repeats and the number of cell states) of the center of mass (CM) of samples randomly selected from each critical state converges to the CM of the whole critical state (y = 0), which shows that the CM of a critical state (see the definition of CM in **section I**) represents its coherent dynamics that emerge in the ensemble of stochastic expression (refer to Fig. 6 in [6]). C) Center of mass (CM) dynamics, which represents coherent dynamics of stochastic expressions, reveals ant-phase between super- and sub-critical states and sequential global perturbations occur at the early - middle 2-cell states (activation → inhibition) and at the middle - late 2-cell states (inhibition → activation). D) Stochastic expressions of the super-critical expression (randomly selected 200 RNAs for mouse embryo) are mixed into stochastic expression of the sub-critical state (4913 RNAs). Euclidian distance of the CM of a randomly selected expression (n) from the mixed ensembles (brown dots) expressions. averaged over 200 repetitions) does not converge to the CM of the sub-critical state, due to the ant-phase coherent behavior of the super-critical state.

**Figure 3** demonstrates distinct coherent-stochastic behaviors emerged in critical states based on i) the law of the large number in each critical state (convergence to its center of mass; **Figure 3B**), ii) different degrees of stochasticity (**Figure 3B**), iii) distinct coherent dynamics (**Figure 3C**) and iv) non-convergence of mixed states into dominant CM (i.e., sub-critical state; **Figure 3D**).

This emergent coherent behavior shows how a single cell can overcome the problem of local stochastic fluctuations in single genes (microscopic rules) by gene expression regulation, which further affirms the statistical significance of self-organization through SOC control of overall expression: genome avalanche and sandpile-type criticality (**Figures 1A and 2**). This collective control in early embryo development points, notably, to the fact that bewildering epigenomic reprogramming molecular processes are regulated by a few hidden control parameters through SOC in terms of collective behaviors even at a single-cell level (see [4] for examples of control at the cell-population level).

Regarding a critical point (CP) in single-cell Th17 cell differentiation and mouse early embryo development, **Figures 1B and 4B,C** suggests that **a single critical point (CP)** may exist in the range of *ln<nrmsf> from* -5.5 to -6.0 for both processes: the onset of the divergence from the scaling occurs at around *ln<nrmsf>* ~ -6.0 for Th17 cell differentiation (**Figures 1B, 4B**), whereas linear regressions in scaling regions for mouse early embryo possesses intersection at around *ln<nrmsf>=* -5.5 ~ -6.0 (a critical point as in **Figures 2, 3A**).

**Figure 4:**
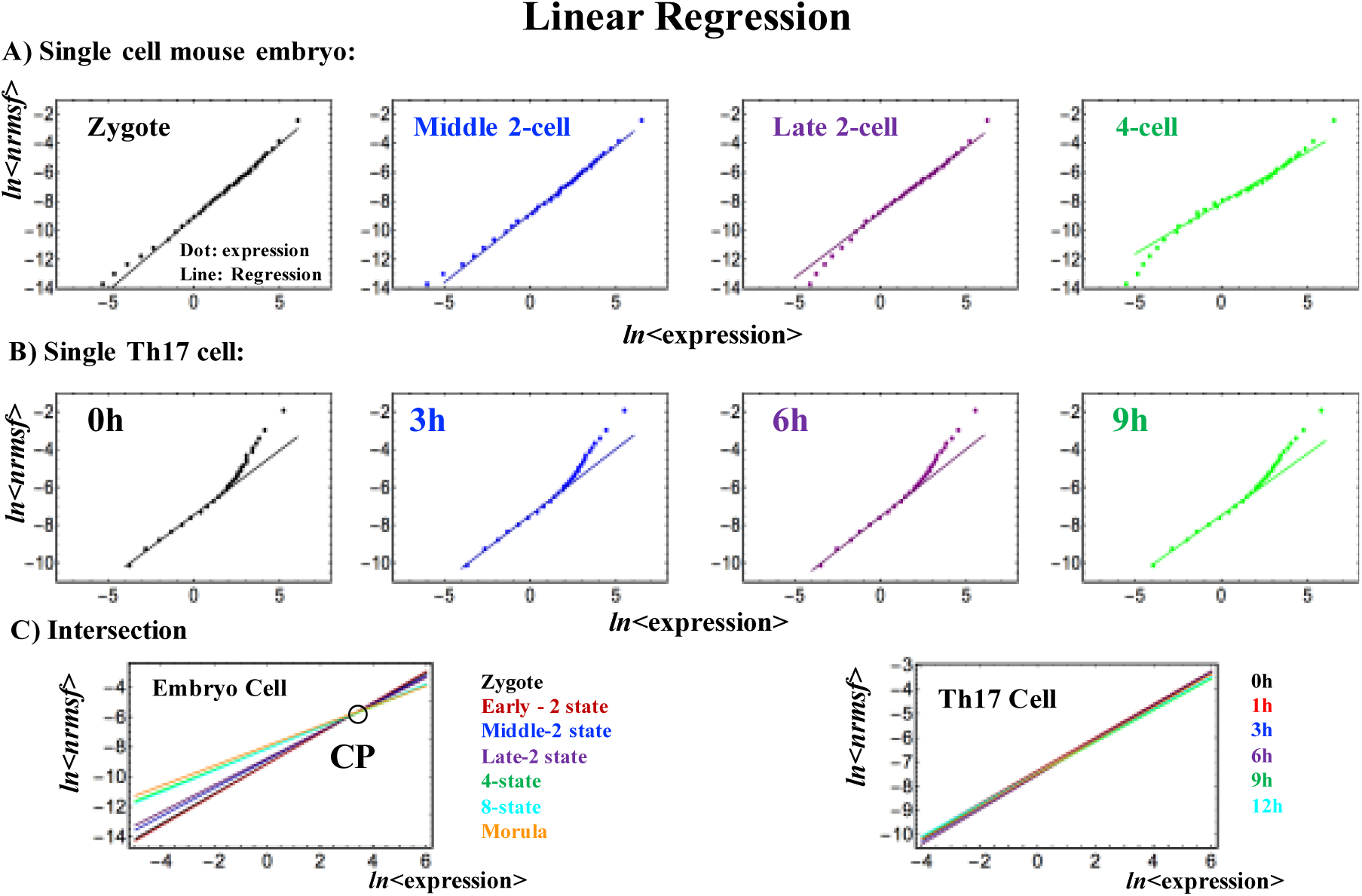
The existence of a fixed critical point in early mouse embryo development: Linear regressions in scaling regions show **A): Linear Regression of ln<expression> vs. ln<*nrmsf*>** (upper row): divergent behaviors occur at low degree of temporal expression for mouse embryo single cell, and **B):** high degree of temporal expression for Th17 single cell, where at around ln<*nrmsf*> ~ -6.0, the onset of the divergence occurs (see also **Figure 1B**, where correlation distance more clearly shows difference in the development of cell states). **C): Intersection of linear regressions** (lower row): Mouse embryo single cell expression intersect around at ln<*nrmsf*>= -5.5 ~ -6.0. This may suggest the existence of near-fixed critical point (CP: **Figures 2, 3A**).

### B) Expression Flux Dynamics Representing the Exchange of Genetic Activity

Here, we elucidate a statistical mechanism to explain how a single mouse embryo cell achieves genome reprogramming by passing through the transition state (**Figure 2**).

**Figure 3B** reveals that the CSB in a critical state corresponds to the scalar dynamics of its CM, *X*(*s*_j_), where *X*(*s*_j_) represents numerical value of a specific critical state (i.e., super-, near- or sub-critical state) at the *j*^th^ cell state, *s*_j_. Thus, the change in the one-dimensional effective force acting on the CM determines the dynamics of *X*(*s*_j_). **Figure 5A** confirms this point, in that the trend of the dynamics of the CM of a critical state follows its effective force (net self-flux dynamics: see the definition below). We now consider that the respective average values of the effective force can serve as baselines, and how perturbation from these baselines occurs dynamically.

It is worth noting here the expression flux between critical states is interpreted as a non-equilibrium system and evaluated in terms of *dynamic network of effective forces*, where interaction flux is driven by effective forces between different critical states is described by second order time difference. From mathematical point of view, the oscillatory phenomenon interpreted with a second-order time difference equation with a single variable is equivalent to inhibitor-activator dynamics given by a couple of first-order time difference equations with two variables.

**Figure 5:**
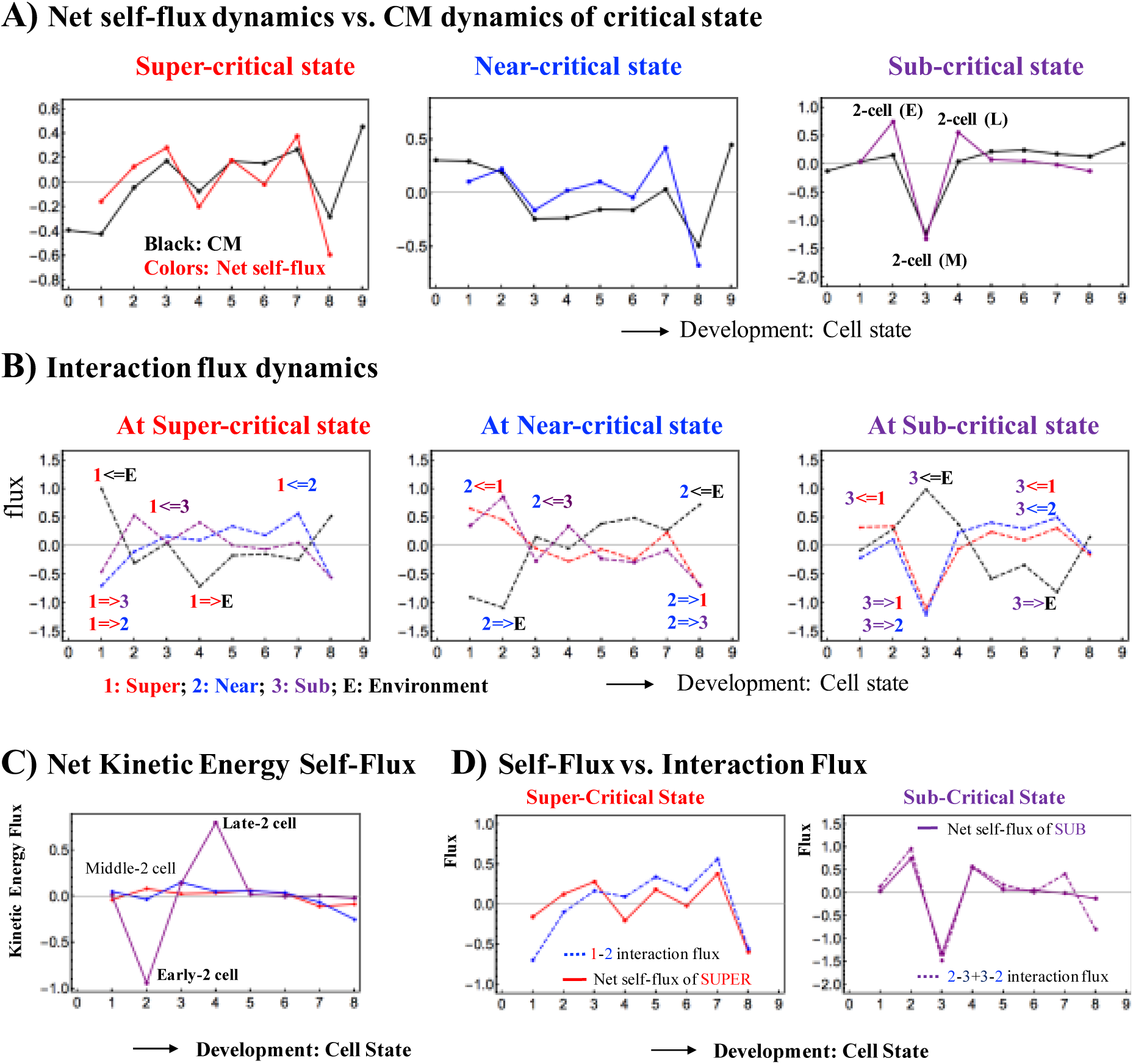
Flux dynamics among interacting critical states: A) The plots show that the net self-flux dynamics (net effective force acting on CM) follow the dynamics of up- or down-regulated CM, where the sign of the net self-flux (i.e., IN and OUT) corresponds to activation (up-regulated flux) for positive responses and inactivation (down-regulated flux) for negative responses. The net self-fluxes of critical states (red: super-critical; blue: near-critical; purple: sub-critical) and the dynamics of the CM of critical states (black lines) are evaluated in terms of time averages. The x-axis represents the development of cell states and the y-axis represents net self-fluxes (black) and their CM dynamics (color). B) The plot shows flux dynamics through crosstalk with the environment. Interaction flux dynamics *i*<=*j* (or *i*=>*j*; color based on that of the *j*^th^ critical state) represent the interaction flux from the *j*^th^ critical state to the i^th^ critical state or vice versa. The results clearly show how global perturbations occur to guide genome reprogramming around the middle 2-cell state (3^rd^ point), based on the activity of the near-critical state and sub-critical state before and after reprogramming (see more in **Figure 7**). C) The change in the net kinetic energy flux from OUT to IN flux (see more in **section II**) shows that sequential global perturbations occur at the early - middle 2-cell states (activation → inhibition) and at the middle - late 2-cell states (inhibition → activation), as in coherent dynamics of critical states (**Figure 3C**). D) The interaction dynamics reveal that the expression dynamics of the sub-critical state (the generator of perturbation) are determined by the net interaction flux between the near- and sub-critical states, and the dynamics of the super-critical state are determined by its interaction flux from the near-critical state (super to near interaction). This image shows how the critical dynamics (temporal change in criticality exists at the boundary of the near- and sub-critical states) affect the entire genome expression system, since the sub-critical state generates autonomous SOC control of overall expression.

The genome is embedded into the intra-nuclear environment, where the expression flux represents the exchange of genetic energy or activity - the effective force produces work, and thus causes a change in the internal energy of critical states. This model shows a statistical thermodynamic picture of self-organized overall expression under environmental dynamic perturbations; the regulation of RNA expression is managed through the mutual interaction between critical states and the external connection with the cell nucleus milieu. The environment here is intended in the broad sense of the baseline gene expression activity adapted to the microenvironment; in thermodynamics terms, the environment can be equated to a ‘thermal bath’.

The effective force can be interpreted as a combination of incoming flux from the past to the present and outgoing flux from the present to the future cell state [4]:

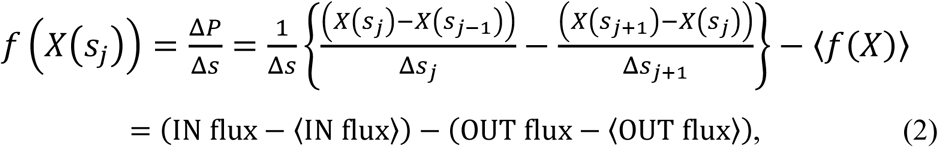

where Δ*P* is the change in momentum with a unit mass (i.e., the impulse: *F*Δ*s* = Δ*P*) and the development of the embryo cell-state is considered as the time-development with an equal time interval (a unit): Δ*s*_j_ = *s*_j_ - *s*_j_ -1= 1 and Δ*s* = *s*_j_+1 - *s*_j_ -1 *=* 2; *s*_j_ corresponds to a specific cell state (such as zygote); the CM of a critical state is 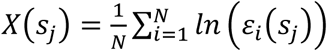 with the natural log of the *i*^th^ expression *ε*_*i*_(*s*), *ln* (ε_*i*_(*s*_j_)) at the *j*^th^ cell state, *s* = *s*_j_ (*N* = the number of RNAs; **Methods**); the average of net self-flux over the number of critical states, <*f*(X)> = <INflux> - <OUTflux>.

The effective force, *f*(*X*(*s*_j_)), is called the *net self-flux of a critical state* at the *j*^th^ cell state *s*_j_. The net self-flux, IN flux - OUT flux, has a positive sign for incoming force (net IN self-flux) and a negative sign for outgoing force (net OUT self-flux). Thus, the CM from its average over all cell states represents up- (down -) regulated expression for the corresponding net IN (OUT) flux.

The *interaction flux* of a critical state, *X* with respect to another critical state or the environment (milieu) *Y* (**Figure 5B**) can be defined as:

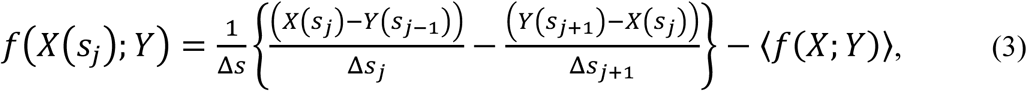

where, again, the first and second terms represent IN flux and OUT flux, respectively, and the net value, IN flux- OUT flux, represents incoming (IN) interaction flux from *Y* for a positive sign and outgoing (OUT) interaction flux to *Y* for a negative sign. *Y* ∈ {Super, Near, Sub, E} where *Y* ≠ *X*: E represents the environment.

Due to the law of force, the net self-flux of a critical state is the sum of the interaction fluxes with other critical states and the environment:

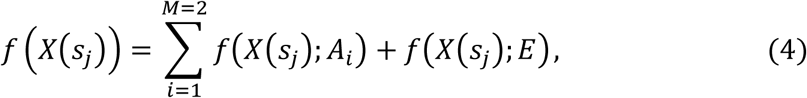

where *f*(*X*(*s*_*j*_); *A*_*i*_) is an interaction flux of *X* with *A*_*i*_ ∈ {Super, Near, Sub} with *A*_*i*_ ≠ *X*, and *M* is the number of internal interactions (*M* = 2), i.e., for a given critical state, there are two internal interactions with other critical states. Equation (4) tells us that the sign of the difference between the net self-flux and the overall contribution from internal critical states, 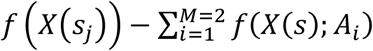, reveals incoming flux (positive) from the environment to a critical state or outgoing flux (negative) from a critical state to the environment.

Regarding average flux, an *average flux balance* exists in terms of the net average fluxes coming in and going out at each critical state (near-zero) through the environment: average expression fluxes: 〈*f*>(*X*)〉 ≈ 0 and 〈*f*(*X*; *y*)〉 ≈0 (Equations (2) and (3)) for the mouse RNA-Seq data (**Methods**). Note: the balance does not hold at each time point/cell state.

This model of expression flux dynamics shows environmental dynamic perturbations in the self-organization of overall expression; the regulation of RNA expression is managed through the mutual interaction between critical states through the cell nucleus.

Under non-equilibrium thermodynamically-open conditions, owe to the break-own of detailed balance, cyclic state-flux among different states emerges as a general property [18]. This is evident in **Figures 6, 7**.

**Figure 6:**
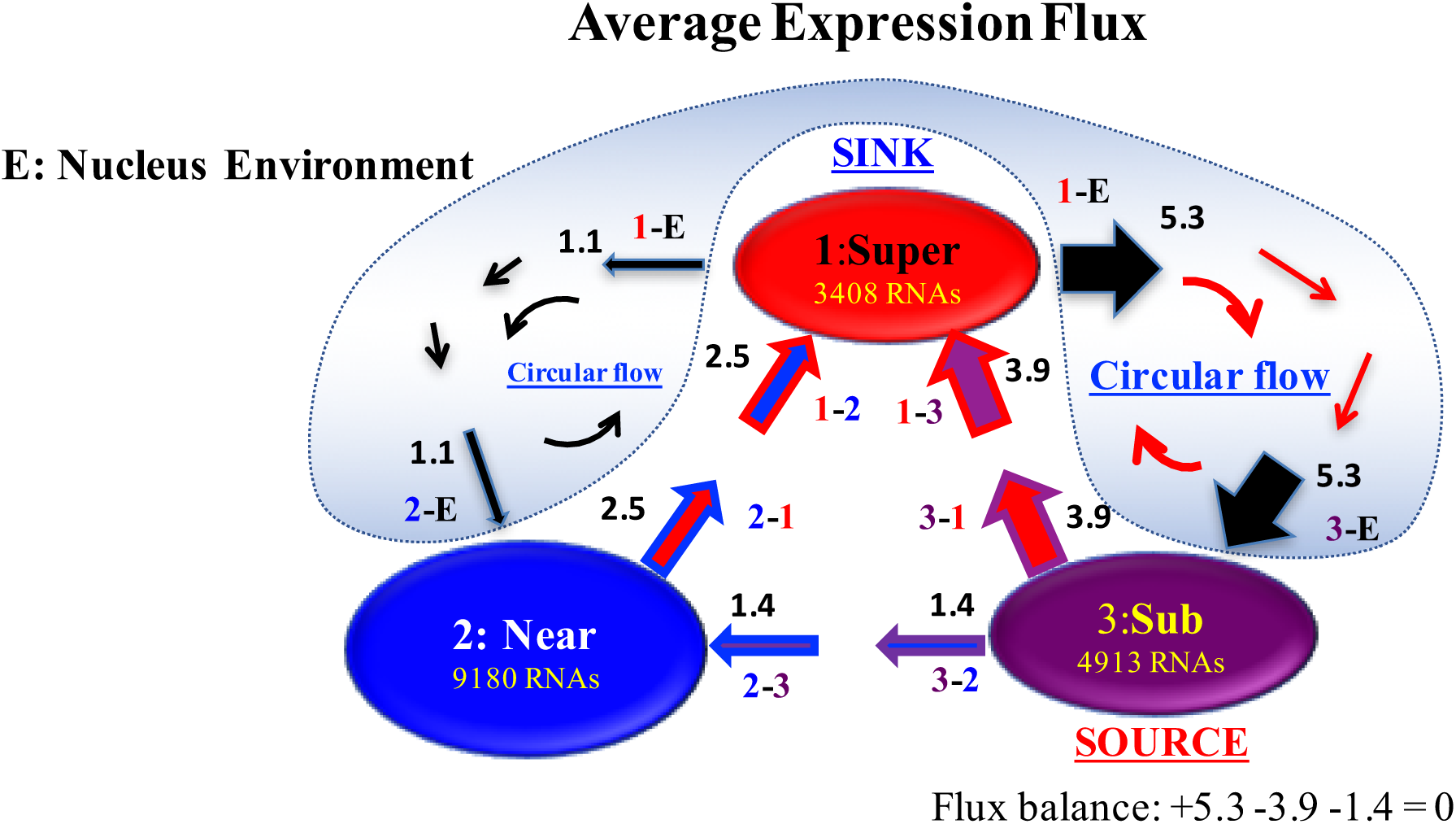
Scheme of SOC-control through the representation of average expression flux in mouse embryo development: Average expression flux network in the mouse embryo genome system reveals that a sub-critical state acts as an internal esource, where IN flux from the environment (shaded blue) is distributed to other critical states. In contrast, a super-critical state acts as an internal esink that receives IN fluxes from other critical states, and the same amount of expression flux is sent to the environment, due to the average flux balance (an example is shown at the sub-critical state). Two cyclic fluxes (Near-Super and Sub-Super) through the environment are seen. Sub-Super cyclic flux forms a dominant flux flow, which generates strong coupling between the super- and sub-critical states accompanied by their anti-phase expression dynamics [6], makes its change oscillatory feedback, and thus sustains autonomous SOC control of overall gene expression. This formation of a dominant cyclic flux with a source and sink provides a universal genome-engine metaphor of SOC control mechanisms, as in terminal cell fates (refer to the Discussion in [4]). Numerics represent average net flux values, where the average net interaction flux of the super-critical state to the environment (black arrow) is decomposed into two parts for the flux-balance at other critical states.

**Figure 7:**
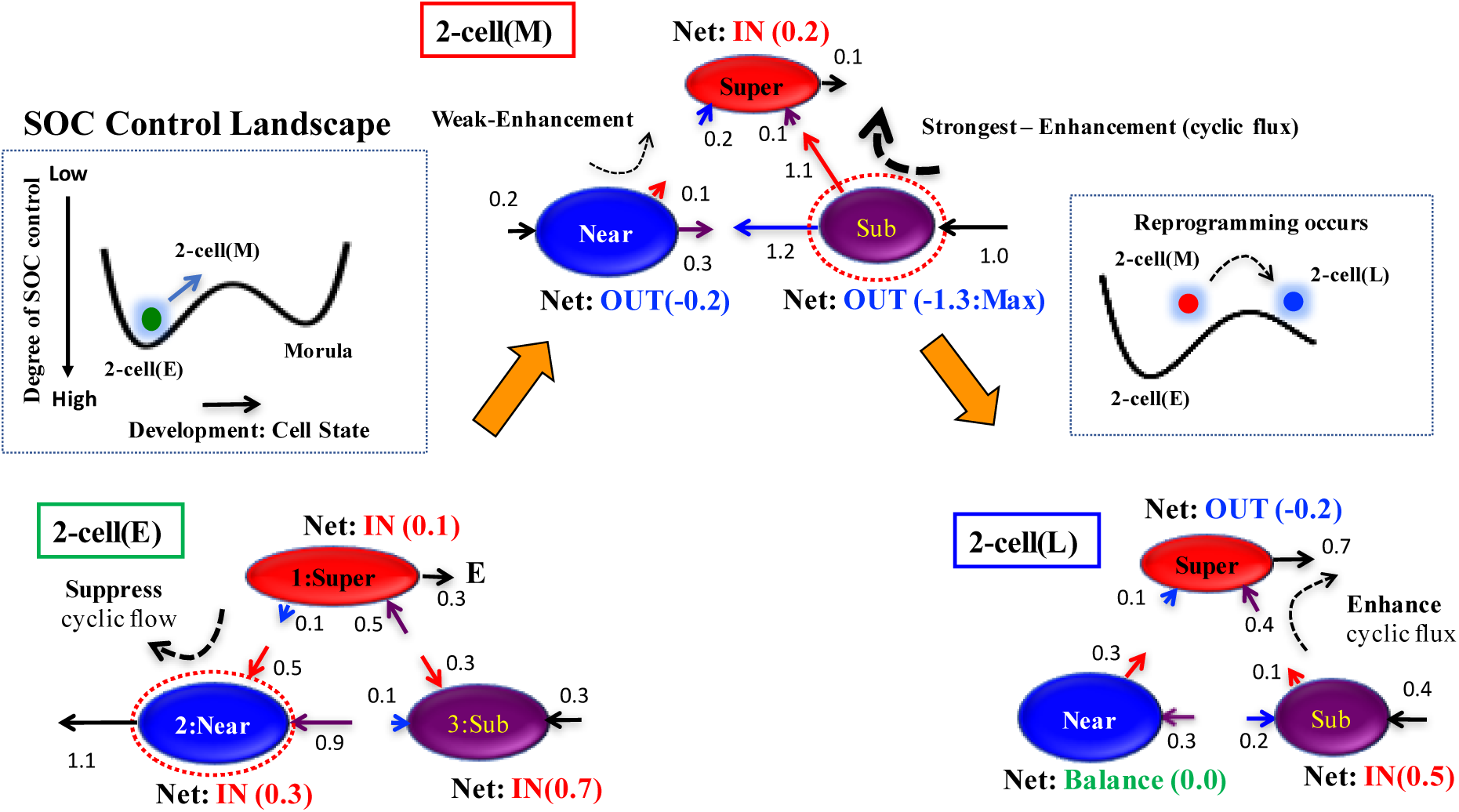
Time-development of SOC control of genome reprogramming revealed through expression flux dynamics: Reprogramming occurs at the middle - late 2-cell states through the transition state of the SOC control landscape (**Figure 2;** more in [4]), which is supported by sequential global perturbations at the early - late 2-cell states (**Figure 3C**). Interaction flux dynamics (**Figure 5B**) reveal a SOC control mechanism of genome reprogramming: i) At the early 2-cell state, interaction flux between Super and Near suppresses cyclic flux. All critical states receive the net IN flux, which indicates that, inside the nucleus, the mouse genome system is activated through the cell milieu (environment), which makes the cell state move up from a valley on the SOC control landscape, as shown in the inset, ii) At the middle 2-cell state, a substantial change in the net interaction flux at critical states occurs from the early 2-cell state to induce a major change in two cyclic fluxes: a change from suppression to the enhancement of cyclic flux (Super - Near), and the strongest enhancement of cyclic flux (Super - Sub). This leads to a reverse change in the net interaction fluxes at the near-critical state (IN to OUT and vice versa) to enhance the cyclic flux (Super - Near) from major suppression in the early 2-cell state. This activation of the sub-critical state (the maximum net OUT flux sending to other critical states) enables the system to pass through the transition state (erasure of zygote criticality: **Figure 2**) to reprogram the mouse embryo genomic state as described in the inset, and iii) In the late 2-cell state, while the enhancement of cyclic flux between super- and sub-critical states becomes weak, the perturbation of average flux activity almost disappears. While the near-critical state becomes balanced, the sub-critical state receives the net expression flux and the super-critical state sends the net flux. i)-iii) show the occurrence of sequential global perturbations to pass through the transition state to achieve single-cell reprogramming. Numerics represent net interaction flux values.

**Figure 6** that shows that how two cyclic fluxes (between Super-Sub critical states, and Super-Near critical states respectively) emerge, while **Figure 5B** reveals broken detailed balance of mutual fluxes between critical states at each time point (see more clearly the broken detailed balance in **Figure 7**), these cyclic fluxes are substantially perturbed before passing the critical transition state (between middle-late 2 cell states; see more in **section II**).

Therefore, it becomes possible to evaluate the temporal change in the open-thermodynamic genetic system, where the expression flux represents the exchange of genetic energy or activity.

### C) Sub-Critical State as a Generator of Perturbation in Genome-Wide Self-Organization

Average net IN and OUT flux flows show how the internal critical states and the external cell nucleus milieu mutually interact (**Figure 6)**. Two cyclic expression flux flows form among critical states: between super- and near-critical states, and *a dominant cyclic expression flux between super- and sub-critical states* in the genomic system.

These cyclic fluxes are considered in light of self-regulatory gene expression by means of a complex epigenetic machinery (methylation processes, long non-coding RNAs, miRNAs, etc): the super-critical state (high variance RNAs) acts as a sink internally to receive genetic information and send it back to the other critical states through the cell environment. On the other hand, the sub-critical state (low-variance RNAs) acts as an internal source of information and sustains (like a generator in an electrical circuit) the interaction with cyclic fluxes.

This implies that the collective behavior of an ensemble of low-variance RNA expression (sub-critical state) plays an essential role in reprogramming in single cells (see a more dynamical discussion below). This dominant cyclic flow also shows that the dynamics of the sub-critical and super-critical states are anti-phase with respect to each other (**Figure 3C**) to form a strong coupling between them [6]. This suggests that the two cyclic fluxes act as feedback flow for the change in criticality to generate coherent oscillatory dynamics of critical states. This dynamic model also provides a mechanism for long-term global RNA oscillation underlying autonomous SOC control generated by the sub-critical state [4,6,19].

The formation of a dominant cyclic flux between a source and a sink provides a genome-engine metaphor for SOC control mechanisms to describe how expression flux is transmitted among critical states: the sub-critical state as a ‘large piston’ for short move and the super-critical state as a ‘small piston’ for large move with an ‘ignition switch’ (near-critical state with a critical point) are connected through a dominant cyclic state flux as a ‘camshaft’, resulted in anti-phase dynamics of two piston movements (refer to Discussion in [4]). This suggests that the genome engine, may be a **Universal mechanism** in the gene expression regulation of mammalian cells.

### D) Change in Criticality: A Global Impact on Whole Genome-Expression System

Here, we further clarify the SOC control mechanism of the reprogramming of single-cell embryo development through the breakdown of initial-state.

The reprogramming occurs at the middle-late 2 cell states: the zygote-state SOC control of overall gene expression (i.e., initial-state global gene expression regulation mechanism) is destroyed through the erasure of the zygote-state criticality. **Figure 5D** shows that the mutual interactions of Sub-Near and Sub-Super determine the net self-flux of the sub- and super-critical states, respectively, representing the effective driving forces acting on their CMs and thus, determine their coherent oscillatory dynamics (**Figure 3C**). This between-states interaction serves as the underlying mechanism of self-regulatory gene expression through the orchestrated cooperation of myriads of epigenetic modifications, transcriptional factors and non-coding RNA regulations to determine the critical-state coherent oscillatory behavior. The essential role played by interactions explains how the temporal change in criticality at the near-critical state, i.e., in expression of the critical gene ensemble of the CP, directly perturbs the sub-critical state (the generator of flux dynamics: **Figure 6**) through their mutual interaction, and the perturbation of this generator can spread over the entire system (refer to Fig 13A in [4]).

### II. SOC Control Mechanism of Genome Reprogramming Through a Critical Transition State

The erasure of initial-state criticality (e.g., in the zygote state) points to the onset of genome reprogramming after the middle 2-cell state. As noted, an initial state can be the early 2-cell state instead of the zygote state; this independent choice of the initial state further confirms the timing of the genome-state change [4].

Interaction flux dynamics (**Figures 5B**, **7**) describes the erasure as a thermodynamical event that passes through a critical transition state, which shows how the genome system can pass through the critical transition state - dynamic perturbation in the average flux in terms of the enhancement-suppression of two cyclic flows around the reprogramming event (**Figure 7**):

i. **Before reprogramming**: In the early 2-cell state, interaction flux (Super - Near) suppresses cyclic flux, where the near-critical state acts as a major suppressor. All critical states receive the net IN flux, which indicates that inside the nucleus, the mouse genome system is activated through the cell milieu (environment).
ii. **Right before reprogramming**: A substantial change in the net interaction flux at critical states occurs beginning in the early 2-cell state and into the middle 2-cell state. These changes induce a major change in the two cyclic fluxes: a change from suppression to the enhancement of cyclic flux (Super - Near), and the strongest enhancement in the dominant cyclic flux (Super - Sub). This leads to the reverse change in the net interaction fluxes at the near-critical state (IN to OUT and vice versa) to enhance the cyclic flux (Super - Near) from the major suppressor at the early 2-cell state. Thus, these reverse cyclic enhancements make the genome system pass through the critical transition state. This interaction model reveals that

a. A major biological event in genome reprogramming through a stochastic pattern is guided by the sub-critical state at the middle 2-cell state, and
b. A detailed open thermodynamic mechanism regarding the erasure of criticality of the early 2-cell state in the middle 2-cell state.
c. The asymmetry becomes the most significant for the IN-and-OUT fluxes between the super- and sub-critical states at the middle 2-cell state, i.e., the greatest time-reversal symmetry breaking is caused. In this respect, it is worth noting that, with no time-reversal symmetry breaking, interaction fluxes between the super- and sub-critical states should be balanced (equal), corresponding to the fundamental framework of detailed balance in equilibrium statistical physics. Whereas, detailed balance should be violated in thermodynamically open system [18].
iii. **After reprogramming:** In the late 2-cell state, while the enhancement of cyclic flux (Super - Sub) becomes weak, the perturbation of average flux activity almost disappears due to passage through the transition state.

Therefore, two major global perturbations, which involve the activation-inhibition of multiple critical states, occur between the early and middle 2-cell states, and between the middle and late 2-cell states during genome reprogramming. This global perturbation event is clearly seen in the net kinetic energy flux [4] in a critical state (**Figure 5C**):

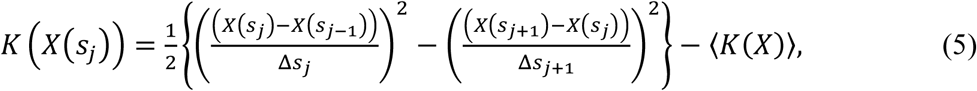

where the kinetic energy of the CM for the critical state with unit mass at *s* = *s^j^* is defined as 1/2. *v*(*s*_*j*_) with average velocity: 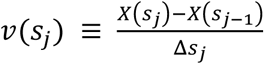.

The above i)-iii) processes around reprogramming can be described in terms of how the genome system can pass through the transition state on the **SOC landscape** (**Figure 3**): the perturbation of self-organization shows a temporal change in the degree of SOC control: a high (low) degree of SOC control points to a weak-local (strong-global) perturbation of the average flux flow. The process that climbs the hill of the *SOC landscape* encompasses the following steps (**Figure 7**): 1). Release from a high degree of SOC control in the early 2-cell state due to the activation of the near-critical state, 2). Passage through the transition state - the lowest degree of SOC control (i.e., non-SOC control) due to the strongest global perturbation at the middle 2-cell state stemmed from the activation of the sub-critical state, which makes SOC control from a high degree to non-control, 3). Return to SOC control in the late 2-cell state.

### III. Discussion and Conclusion

Thermodynamics allows for incredibly precise predictions thanks to the ‘generality of its premises’ (Einstein’s words [20]), this allowed us to grasp the essentials by skipping the (largely unknown) detailed biological mechanisms and focusing on the phenomenology of the genome expression global changes. The erasure of epigenetic marks (refer to the Discussion in [4]) is consistent with thermodynamics perspective beside its actual mechanism.

The time-development of sandpile criticality reveals the existence of a transition state which in turn suggests that programming of mouse embryo occurs through the transition state. As a proof of this concept, it was demonstrated that EGF-stimulated MCF-7 cells do not erase sandpile criticality (see Fig. 5A in [4]), i.e., no genome-state change (consistent with the experiment no cell-differentiation occurs [21]). This shows that the event of single-cell programming is related to overcoming of the transition state [22,23], which is a typical event in thermodynamic reaction mechanism. Moreover, the timing of the reprogramming does not depend on the selection of an initial cell state. The result helps us to obtain a quantitative appreciation of the still largely qualitative notion of the epigenetic landscape.

Intriguingly, our statistical thermodynamics approach reveals that the collective behavior of an ensemble of low-variance RNA expression (sub-critical state), which shows only marginal changes in expression and consequently are considered to be devoid of any interest, guides the genome to pass through the transition state (erasure of initial-state sandpile-type criticality) to reprogram the mouse embryo. The sub-critical state gene ensembles act as a driving force to transmit their potentiality, or energy of coherent transcription fluctuations, to high-variance genes (the genome-engine mechanism [4]). We observed also this generator role of the sub-critical state in cell differentiation [4] (MCF-7 human cancer cells and HL-60 human promyelocytic leukemia cells). Thus, genome-engine mechanism may provide a universal SOC control mechanism in the genome system. 20

Sandpile-type critical point (CP) exists around the edge between the near- and super-critical states in the mouse genome. The erasure of sandpile-type criticality through embryo development induces dynamic change in interaction flux dynamics between near- and sub-critical states, which determines the activation-inhibition dynamics of the sub-critical state. The sub-critical state generates autonomous SOC control of overall expression, and thus the critical dynamics (temporal change in criticality) affect the entire genome expression system. This suggests that a critical gene ensemble of sandpile-type criticality (i.e., critical point) should exist to affect the entire genome expression. Therefore, elucidation of the molecular mechanism that guides the genome system through a transition state is expected to unveil molecular clues as to how a single cell can succeed or fail at reprogramming, such as in an iPS single cell.

The establishment of a material basis for the observed phenomenology by unveiling the molecular mechanism on the criticality of gene ensemble is awaited as the next study, which may lead us toward a comprehensive understanding of single-cell reprogramming.

## IV. Methods

### Biological Data Sets

We analyzed the following mammalian RNA-Seq data:

i) Early embryonic development in human and mouse developmental stages in RPKM (Reads Per Kilobase Mapped) values; GEO ID: GSE36552 (human: *N* = 20286 RNAs) and GEO ID: GSE45719 (mouse: *N* = 22957 RNAs), which have 7 and 10 embryonic developmental stages (experimental details in [24] and [16], respectively):

Human: oocyte (m=3), zygote (m=3), 2-cell (m=6), 4-cell (m=12), 8-cell (m=20), morula (m=16) and blastocyst (m=30),

Mouse: zygote (*m*=4), early 2-cell (*m*=8), middle 2-cell (*m*=12), late 2-cell (*m*=10), 4-cell (*m*=14), 8-cell (*m*=28), morula (*m*=50), early blastocyst (*m*=43), middle blastocyst (*m*=60) and late blastocyst (*m*=30), where *m* is the total number of single cells.

ii) T helper 17 cell differentiation from mouse naive CD4+ T cells in RPKM values, where Th17 cells are cultured with anti-IL4, anti-IFNγ, IL-6 and TGF-β, and Th0 cultures provide control cells that receive TCR activation in the absence of exogenous polarizing cytokines (IL-6 + TGF-β) (details in [25]); GEO ID: GSE40918 (mouse: *N* = 22281 RNAs), which has 9 time 21 points: *t*0 = 0, *t*1 = 1,3,6,9,12,16,24, *tT*=8 = 48h for Th17, and 6 time points: *t*0 = 0, *t*1 = 1,3,6,16, *tT*=5 = 48h for Th0.

RNAs that had RPKM values of 0 over all of the cell states were excluded. Random real numbers in the interval [0-1] generated from a uniform distribution were added to all expression values in the analysis of sandpile criticality (**Figure 2**; no addition of random numbers in the rest of the Figures). This procedure avoids the divergence of zero values in the logarithm. The robust sandpile-type criticality through the grouping of expression was checked by multiplying the random number by a positive constant, *a* (*a*< 10), and we set *a* = 0.01. Note: The addition of large random noise (*a*>>10) destroys the sandpile CP.

### Normalized Root Mean Square Fluctuation (*nrmsf*)

*Nrmsf* is defined by dividing *rmsf* (root mean square fluctuation) by the maximum of overall {*rmsf_i_*}:

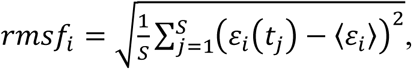

where *rmsfi* is the *rmsf* value of the *i*^*th*^ RNA expression, which is expressed as *εi*(*s^j^*) at a specific cell state *s^j^* (in mouse, *S* = 10: *s*1 *=* zygote, early 2-cell, middle 2-cell, late 2-cell, 4-cell, 8-cell, morula, early blastocyst, middle blastocyst and *s*10 *=* late blastocyst), and 〈*ε*_*j*_〉 is its expression average over the number of cell states. Note: *nrmsf* is time-independent variable and an order parameter for self-organization of genome expression as demonstrated in our previous works [4-6].

### Bimodality Coefficient

Sarle’s bimodality coefficient for a finite sample (*b*) [26] is given by

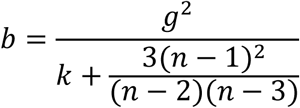

where *n* is the number of items in the sample, *g* is the sample skewness and k is the sample excess kurtosis.

It is worth noting that gene expression in single cell RNA-Seq data we used in the present paper do have many zero values. This is not an artifact but a consequence of the ‘toggle-switch’ mechanism of gene regulation [27] and are essential for the understanding of regulation mechanisms. On a purely computational perspective, zeroes alter the ‘standard values’ of the bimodality coefficient (*b*= 5/9 ~ 0.56 indicates a bimodal or multimodal transition from a unimodal profile), this implies we must limit ourselves to consider bimodality coefficient a step-like behavior as the signature of a transition (**Figure 3A**) without entering in the details of their actual values.

### SOC Control Mechanism of Overall Expression

A self-organized critical transition (SOC) in whole-genome expression plays an essential role in the change of the genome expression state - SOC control of overall expression at both the population and single-cell levels [4-6]. The basic findings of these studies can be summarized as:

i). SOC of overall expression does not correspond to a phase transition from one critical state to another. Instead, it represents self-organization of the coexisting critical states through a critical transition, i.e., SOC consolidates critical states into a genome expression system (called e*SOC control of overall expression*) accordingly to temporal expression variance (*nrmsf*). *Nrmsf* acts as an order parameter in self-organization. In critical states, distinct coherent (collective) behaviors emerge in ensembles of stochastic expression (coherent-stochastic behavior), where coherent dynamics of high-variance gene expression (super-critical state) is anti-phase to that of low-variance gene expression (sub-critical state).

ii). The characteristics of the self-organization through SOC become apparent only in the collective behaviors of groups with an average of more than around 50 genes (mean-field approach). The same value of 50 genes (or around this value) as the threshold for the onset of coherent ensemble behavior was previously recognized in a completely different context and by different analytical techniques [6,8,9]. This effect is clearly a statistical one, but not a esimple statistical one in the sense it is a *proxy* of the underlying gene regulation network (GRN).

iii). Self-organization occurs through distinguished critical behaviors: *sandpile-type criticality* and *scaling-divergent behavior*:

a. Sandpile-type critical behavior (criticality) based on the grouping of expression according to the fold change in expression. A summit of the sandpile represents the critical point (CP); as the distance from the CP increases, the divergence of two different regulatory behaviors occurs, which represent up-regulation and down - regulation, respectively. Furthermore, in the vicinity of the CP according to *nrmsf*, in terms of coherent expression, self-similar bifurcation of overall expression occurs to show a unimodal-bimodal transition (Figures. 1, 3A in [6]: MCF-7 cancer cells) and a step function like transition (Figure 3A in [4]: HL-60 cancer cells); these indicates the existence of cell-type specific transitions. Thus, since a critical behavior and a critical transition occur at the CP, we can characterize it as a *sandpile-type transition.*
b. Scaling-divergent behavior (genomic avalanche) based on the grouping of expression according to *nrmsf*: a nonlinear correlation trend between the ensemble averages of *nrmsf* and gene expression at each time point, which has both linear (scaling) and divergent domains in a log-log plot; the onset of divergence occurs at the CP: order (scaling) and disorder (divergence) are balanced at the CP (), which presents a genomic avalanche. The scaling-divergent behavior reflects the co-existence of distinct response domains (critical states) in overall expression. *Distinct critical behaviors from different averaging behaviors occur (numerically) at around the same CP*.

The occurrence of a temporal change in criticality directly affects self-organization in the entire genomic system (Figure 13 in [4]). The genome-state change (cell-fate change in the genome) occurs in such a way that the initial-state SOC control of overall gene expression - initial-state global gene expression regulation mechanism is destroyed through the erasure of an initial-state criticality.

## Acknowledgements

MT sincerely thanks the Institute for Advanced Biosciences, Keio University, Tsuruoka City, the Yamagata prefectural government, Japan, SEIKO Research Institute (SRI) for Education, Japan, and Mr. Fumiaki Kikuchi for allowing him to complete this research project at Keio University. The authors are thankful to Dr. Midori Hashimoto for preparing Figures, and Drs. Jekaterina Erenpreisa and Mesut Tez for fruitful and critical discussions.

